# Involvement of mechanical cues in the migration of Cajal-Retzius cells in the marginal zone during neocortical development

**DOI:** 10.1101/2021.10.25.465770

**Authors:** Ana López-Mengual, Miriam Segura-Feliu, Raimon Sunyer, Héctor Sanz-Fraile, Jorge Otero, Francina Mesquida-Veny, Vanessa Gil, Arnau Hervera, Isidre Ferrer, Jordi Ortín, Jordi Soriano, Xavier Trepat, Ramon Farré, Daniel Navajas, José A. del Río

## Abstract

Emerging evidence points to coordinated action of chemical and mechanical cues during brain development. At early stages of neocortical development, angiogenic factors and chemokines such as CXCL12, ephrins, and semaphorins assume crucial roles in orchestrating neuronal migration and axon elongation of postmitotic neurons. Here we explore the intrinsic mechanical properties of the developing marginal zone of the pallium in the migratory pathways and brain distribution of the pioneer Cajal-Retzius cells. These pioneer neurons are generated in several proliferative regions in the developing brain (e.g., the cortical hem and the pallial subpallial boundary) and migrate tangentially in the preplate/marginal zone covering the upper portion of the neocortex. These cells play crucial roles in correct neocortical layer formation by secreting several molecules such as Reelin. Our results indicate that the motogenic properties of Cajal-Retzius cells and their perinatal distribution in the marginal zone are also modulated by both chemical and mechanical factors, by the specific mechanical properties of Cajal-Retzius cells, and by the differential stiffness of the migratory routes. Indeed, cells originating in the cortical hem display higher migratory capacities than those generated in the pallial subpallial boundary which may be involved in the differential distribution of these cells in the dorsal-lateral axis in the developing marginal zone.

## Introduction

Cajal-Retzius (CR) cells were first described by Santiago Ramón y Cajal and Gustaf Retzius (in 1890 and 1892, respectively) [1, 2]. These cells are early-generated neurons located in marginal zone/cortical layer I that split from the embryonic preplate to form the marginal zone when the cortical plate develops during early cortical development (e.g., see [2–6]). Although several differences in CR cell phenotype, marker, physiology, and fate have been described in different mammals (e.g. [7–9]), Reelin expressed by mouse CR cells during cortical development modulates the appropriate migration of cortical plate neurons, actively participating in neuronal network activity in developing marginal zone/layer I (e.g., [3]). In rodents, CR cells have the capacity to generate action potentials, establishing synaptic contacts in the marginal zone/layer I and receiving excitatory and GABAergic and non-GABAergic inputs [3, 4, 10–15]. Mouse CR cells are mainly generated in three neurogenic areas: the cortical hem (CH) [16, 17], the septum retrobulbar area (SR), and the pallial subpallial boundary (PSB) [18]. Shortly after generation, CR cells migrate through the preplate/marginal zone to populate the entire cortical surface following specific rostro-caudal and latero-tangential processes [16, 18–23]. This dorsal-ventral migration of CR cells as well as subplate neurons thorough the preplate has been reported to play a crucial role in regionally defining the developing neocortex [24]. Birthdates of cortical CR cells are between embryonic days 8.5 and 13.5 (E8.5- 13.5) in the mouse, with a maximum between E9.5 and E12.5 [25–27], although a recent study points to a supply of CR cells from the olfactory bulb at protracted embryonic stages [28]. During the first and second postnatal week, mouse CR cells disappear from layer I by programmed cell death [26, 29]. In fact, both their distribution and their differential disappearance play a role in neocortical regionalization and maturation [30, 31].

Genetic screening of CR cells has revealed that a large number of factors are involved in their generation, migration, and maturation, such as *p73*, *p21*, *Zic1-3, Lhx5 and Fgf8*, *Tbr1 and 2*, *MDGA1*, *Emx1* and *Emx2*, *Nectin1*, *Dmrt*, *Dbx1*, *Foxg1*, *Ebf2, Foxc1, LIM-homeobox* genes, and *miRNA9,* among others [25, 32–44]. Concerning migration, several molecules have been identified as regulators of CR-cell migration and distribution in the marginal zone, e.g., CXCL12, Eph/ephrin, and Pax6 [22, 45–48]. In fact, CXCL12 (also termed stromal derived factor I, SDF-1), secreted by meningeal cells, is considered to be mainly responsible for hem-derived CR-cell migration through CXCR4-CXCR7 receptors expressed in CR cells. Surprisingly, the migration at the subpial position of CR cells in *CXCR4-/-, CXCR7-/-,* and *CXCL12-/-* mice, although affected, is largely maintained at dorsal pallial levels [49]. This is in contrast to other studies displaying relevant changes in CR-cell location and cortical layering after the chemical removal of meningeal cells or genetic modification, suggesting that other factors associated with meninges are involved in CR-cell migration and distribution [45, 50, 51]. Angiogenic factors present in the marginal blood vessels associated with meninges such as VEGF, Sema3E, and Ephrins have emerged as important cellular cues regulating the migration of CR cells, as is the case in other developmental processes [52–54].

In addition, evidence emerging from recent research shows that, in parallel to chemical cues, neural morphogenesis, neuronal migration, and axon navigation are processes also governed by sensing the mechanical properties of the extracellular milieu (e.g., Young’s modulus and topography) and neighboring cells during development [55–58]. These interactions influence the maturation and differentiation of particular neurons based on transduction of those external mechanical forces into intracellular biochemical signaling via a mechanical- transduction process [58, 59]. This mechanical-transduction process involves the action of integrins and other elements linking extracellular matrix (ECM) to cellular cytoskeleton dynamics [60]. In addition, specific signaling mechanisms such as mechanosensory receptors (e.g., Piezo) and Hippo/YAP pathways are players in the mechanical-transduction process in several cell types and tissues (e.g., [61, 62]), including neural tissue (e.g., [63]). Considering matrix stiffness during mouse cortical development, Iwasita and coworkers measured, by means of atomic force microscopy (AFM), the values for Young’s/Elastic modulus (*E*) of the cortical plate (CP), the intermediate zone (IZ), the subventricular zone (SVZ), and the ventricular zone (VZ) in coronal sections of the prospective parietal cortex of the mouse at different postnatal stages (from E12.5 to E18.5) [64]. In a broad sense, all values (for all cortical layers) increased from E12.5 with a peak at E16.5 and then decreasing (see Figs. 1-2 in [64]) at late (E18.5) embryonic stages. Young’s modulus values ranked (for the CP) from 30.1 Pa at E12.5 to a maximum of 108.4 Pa (E16.5), decreasing to 57.4 Pa at E18.5. However, preplate/molecular layer/layer I was not thoroughly analyzed in the study.

**Fig. 1.**
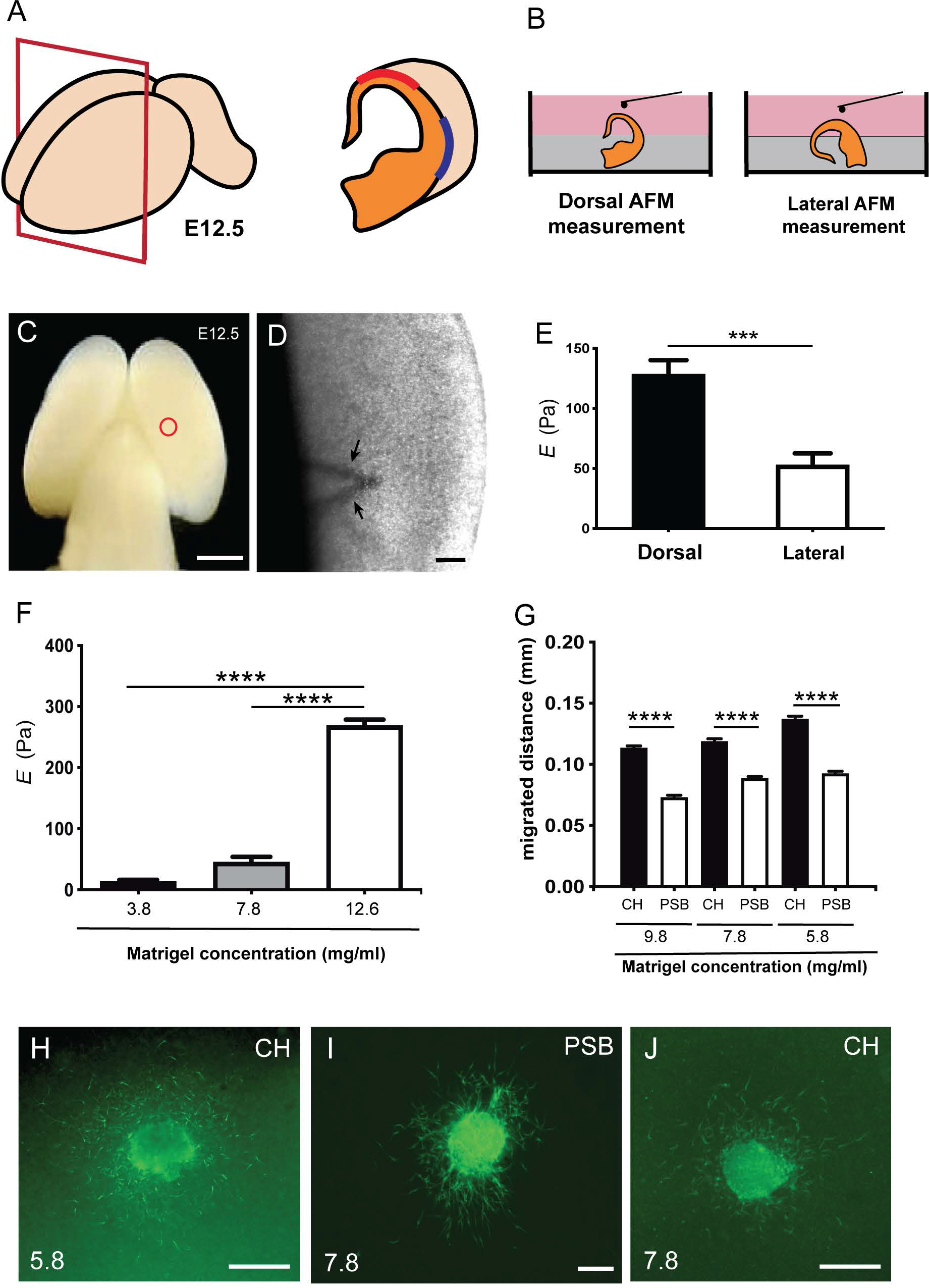
Differential stiffness between dorsal and lateral regions of marginal zone in developing mouse embryos. **A-B)** Scheme illustrating the procedure of placement of the telencephalic hemispheres of the embryo (E12.5), embedded in agarose to obtain dorsal and lateral measurement using the BIO-AFM. **C-D)** Scheme (C) and low magnification photograph (D) obtained from the BIO-AFM illustrating the location of the V-shaped cantilever (circle in C and arrows in D) in the surface of the marginal zone. **E)** Histogram showing the results of the BIO- AFM experiments; *E* values are displayed in the y axis in Pa. **F)** Rheometric values obtained after the analysis of three different hydrogels. The concentration of the total protein of the analyzed hydrogels is shown in the x axis. **G)** Bar plots comparing the amount of differential migration of CR cells (obtained from CH or PSB) for gradually higher matrigel concentrations. **H-J)** Examples of CH (H-J) and PSB (I) cultured explants in different hydrogel concentrations (5.8 and 7.8 mg/ml) immunostained against CALR to identify CR cells. CH: cortical hem; PSB: pallium subpallium boundary. Data in E, F, and G are presented as mean ± s.e.m.; *** *p* < 0.001 and **** *p* < 0.0001. Scale bars C = 1 mm, D = 500 μm, H and J = 300 μm and I = 300 μm.

**Fig. 2.**
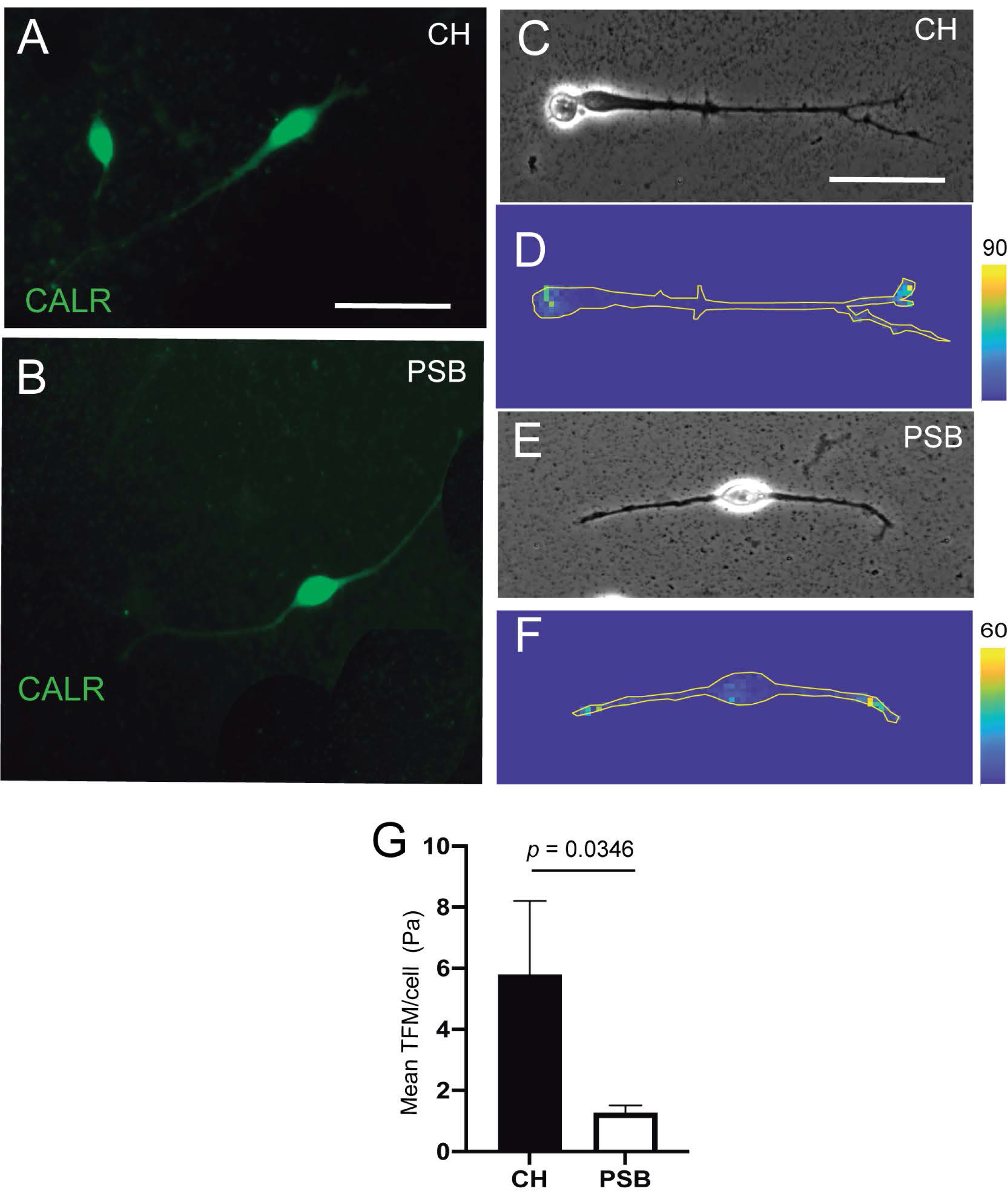
TFM measurements of CH and PSB-derived CR cells in PAA gels. **A-B)** Examples of CR cells stained using *α*-CALR antibodies derived from CH (A) and PSB (B). **C-F)** Phase contrast (C and E) and constraint force maps (D and F) of CR cells derived from CH (C-D) and PSB (E-F) after TFM analysis. Forces triggered by CR cells are displayed with different color following the color scale shown in D and F. **G)** Histogram showing the results of the TFM analysis. Plotted values represent the mean of the forces generated by the different CR cells. Note that the values in the histogram represented are the means of the analyzed forces divided by the total pixels occupied by the analyzed CR cell. The shape of the cell scan be seen in the constraint images. In addition, note that the maximum values of traction are morphologically located in the tips of their neurites in CH- and PSB-derived CR cells (D and F). CH: cortical hem; PSB: pallium subpallium boundary. Data in G are presented as mean ± s.e.m. and the *P* value indicated in G was obtained using the permutation test (one-tailed) (see Material and Methods for details). Scale bars, A = 50 μm pertains to B, C = 50 μm pertains to D-F.

Concerning CR cell migration, a pioneer study described differences in the migratory properties of these cells depending on their origin (rostral *vs* medial) after ectopic transplantation in different areas of the embryonic cortical hem [48].

Thus, the study revealed, for the first time, that the local environment in parallel to guidance molecules can modulate the migration of CH-derived CR cells [48]. In addition, Barber *et al.* demonstrated differences in migratory speed of subsets of CR cells [65]. In fact, using *in vitro* experiments of the complete pallium, Barber et al. described how SR-derived and CH-derived CR cells migrate and then stop their migration at the dorsal pallium levels; whereas PSB derived CR cells migrate in the rostral caudal axis in the lateral part of the pallium [65].

In this study, we aimed to explore whether putative differences in the mechanical properties between dorsal and lateral parts of the developing marginal zone/layer I also influenced the migration and distribution of CR cells derived from the CH and the PSB of the mouse. Our data reveal that both the stiffness differences between medial and lateral regions of the marginal zone as well as the intrinsic mechanical properties of CR cells contribute to their migration from the CH and PSB in the dorsal and lateral parts of the developing cortex.

## Results

### Dorsal to lateral stiffness (*E*) differences are present in developing marginal zone of developing mouse cortex

As a first set of experiments we developed BIO-AFM measurements in dorsal and lateral parts of the cortical surface of embryonic mice (E12.5) (Fig. 1). At this embryonic stage, a lateral growth of the pallium takes place [66] and CR cells tangentially migrate through the marginal zone and cover the entire pallial surface (see introduction for references). In our experiments we focused on the dorsal and lateral portions of the upper cortical surface (Fig. 1B-D). Our BIO-AFM results indicate clear differences in stiffness between the dorsal and lateral portions of the neocortical surface (dorsal: 128.1 ± 12.08 Pa *vs* lateral: 52.46 ± 12.08 Pa, mean ± s.e.m., *** *p* = 0.0002) (Fig. 1E) (n = 9 for each condition).

### Differential migration of CH and PSB-derived CR cells in different Matrigel^TM^ concentrations

Next, we aimed to explore whether these stiffness differences might affect the migration of CR cells (Fig. 1F-J). Classical studies analyzing CR cell migration used 3D-Matrigel^TM^ hydrogels as a migration substrate (e.g., [46, 53]). In order to determine whether the stiffness of the environment could modulate in the migration of CR cells in a region-specific manner, we first analyzed the stiffness of different Matrigel^TM^ concentrations using rheometric analysis (Fig. 1F). We selected the Matrigel^TM^ concentrations taking into account the total protein level in the different batches obtained from the supplier (see Materials and Methods for details) in terms of the generation of a homogenous hydrogel at each of these concentrations (see [67] for examples). Data illustrate that, as expected, high Matrigel^TM^ concentrations led to significantly higher elastic moduli *E* (Fig. 1F), with *E* decreasing from 12.6 mg/ml to 3.8 mg/ml Matrigel^TM^ dilutions. Measured *E* at 1 Hz for 12.6 mg/ml was 268 ± 10.11 Pa (n = 6), for 7.8 mg/ml it was 45.39 ± 8.93 Pa (n = 5), and for 3.8 mg/ml it was 13.34 ± 3.02 Pa (n = 4), data as mean ± s.e.m. (Fig. 1F). When comparing the elasticity of the hydrogels with those previously obtained in BIO-AFM, *E* values obtained using 12.6 mg/ml of Matrigel^TM^ were around 2 times than those measured in the *in vivo* BIO-AFM. However, the data obtained using 7.8 mg/ml were similar to those observed in lateral regions of the apical surface and the values using 3.8 mg/ml were 4 times lower than the lateral telencephalic portion of the marginal zone. For this reason, we used three different concentrations of Matrigel^TM^ ranging around physiological values as measured: 5.8 mg/ml, 7.8 mg/ml and 9.8 mg/ml.

As indicated in several studies, CR cells derived from the CH and PSB showed an almost non-overlapping distribution in the developing marginal zone-layer I (e.g., [68]). Taking this into account, we cultured CH and PSB explants in Matrigel^TM^ hydrogels with different concentrations. CR cells from CH were able to migrate longer distances in 5.8 mg/ml, 7.8 mg/ml, and 9.8 mg/ml Matrigel^TM^ dilutions (Values for 9.8 mg/ml: CH = 113.4 ± 1.7 μm (n = 1632) *vs* PSB = 72.86 ± 1.3 μm (n = 915). Values for 7.8 mg/ml: CH = 131.7 ± 2.1 μm (n = 1471) *vs* PSB = 88.52 ± 1.4 μm (n = 1911). Values for 5.8 mg/ml: CH = 137.1 ± 2.2 μm (n = 1737) *vs* PSB = 92.35 ± 2.2 μm (n = 1194); all mean ± s.e.m.) (Fig. 1G and two supplementary movies S1 and S2). This suggests that i) hem-derived CR cells have greater motogenic capacity than PSB-derived cells when migrating in hydrogels with the same *E* value, and ii) hem-derived CR cells are able to migrate greater distances in hydrogels displaying *E* values closer to those observed in the dorsal portion of the pallium in contrast to PSB-derived CR cells. In Fig. 1H-J we offer some examples of the distribution of CALR-positive CR cells after completing their migration for 2 DIV in different Matrigel^TM^ concentrations.

### Hem-derived CR cells displayed greater mechanical forces than PSB-derived CR cells in polyacrylamide gels

In a next set of experiments, we aimed to determine whether these migratory differences could be attributed to intrinsic differences between CH-derived or PSB-derived CR cells in order to generate mechanical forces when cultured on polyacrylamide (PAA) substrates. First, we generated PAA gels with very low *E* using the protocol published in [69, 70]. We obtained soft PAA gels with *E* values around **≍** 40-100 Pa. This *E* value is the lowest stiffness that allowed us to obtain good distribution of the nanoparticles used in traction force microscopy (TFM). Below these values the PAA is not stable and does not generate reliable TFM measurements. We aimed to analyze the behavior of CH- and PSB-derived CR cells when cultured on these low-Pa PAA gels. CR cells adhered to the PAA gel and did not migrate but generated forces on the substrate (Fig. 2). In some cases, due to the absence of a 3D-hydrogel environment, CR cells modify their morphology from the typical unipolar to a more bipolar shape as also observed in other studies in 2D-cultures [22]. In previous experiments we also categorized the cell morphologies as CR cells in PAA gels using CALR immunostaining (Fig. 2A-B). Thus, we developed the TFM measures in isolated CR cells with these morphologies (Fig. 2C and F) and did not analyze TFM in cells with multipolar morphology nor did we group them to compare equal populations. TFM results demonstrated, as expected, that CR cells independently of their origin, do not generate large forces to the PAA substrates compared to other cell types. TFM analysis reported that CH-derived CR cells can develop greater forces on the substrate when compared to PSB-derived ones (*p* = 0.0346, one tail permutation test, n = 18 and 14, respectively) (Fig. 2G). This also points to differing intrinsic mechanical properties between CH and PSB-derived CR cells that might allow CR cells to sense the different stiffness of the marginal zone. One potential mechanism to sense stiffness is the expression of mechanosensory channels [71]. To assess this, we developed a loss of function experiment using the mechanosensory channel inhibitor GsMTx-4 [72] (Fig. 3). This compound is a spider venom that inhibits cationic mechanosensitive channels. In fact, although the specific mechanisms of the drug have not been fully determined, when GsMTx-4 is applied to several cell types expressing mechanosensitive channels (e.g., Piezo channels) they remain open and are able to increase intracellular Ca^2+^ level [73]. Taking this into account, we cultured hem-derived explants, and after 40 hours in order to obtain isolated CR cells, cultures were incubated with Fluo4-AM. After incubation, the changes in the Ca^2+^ waves in CR cells were analyzed using NETCAL Software [74]. First, we checked the health status of cultured CR cells in 1% methylcellulose containing medium by analyzing their depolarization, using KCl (Fig. 3A-C). After KCl treatment, an increase in the fluorescence *ΔF/F0* values was observed in all analyzed CR cells (Fig. 3C). Next, we developed similar experiments, first incubating CR cells with the inhibitor GsMTx-4 and then after that with KCl (Fig. 3D-E). Results demonstrated that treatment with GsMTx-4 transiently increased intracellular calcium levels in CR cells, reducing their migration (CH, Veh = 130, 8 ± 2.5; GsMTx-4 = 124, 4 ± 3.5, mean ± s.e.m., ** *p* = 0.0015, n = 1218 and 782, respectively) (Fig. 3E-F). For PSB, CR-cell migration was lower after incubation with the inhibitor but did not reach statistical significance (*p* = 0.063, n = 1289 for GsMTx-4 and 382 for Veh) (Fig. 3F). In addition, we analyzed whether the blockage of cytoskeleton proteins and myosin II also impaired their migration. These experiments showed, as expected, that inhibiting tubulin (Nocodazole; *p* < 0.001, n = 21) and myosin II (Blebbistatin; *p* = 0.032, n = 22) almost blocked the migration of CR cells (Fig. 3G). Taken together, the present data demonstrate that CR cells can generate mechanical forces to the substrate (CH > PSB-derived CR cells), but that cytoskeletal disruption impairs their migration on Matrigel^TM^ hydrogels.

**Fig. 3.**
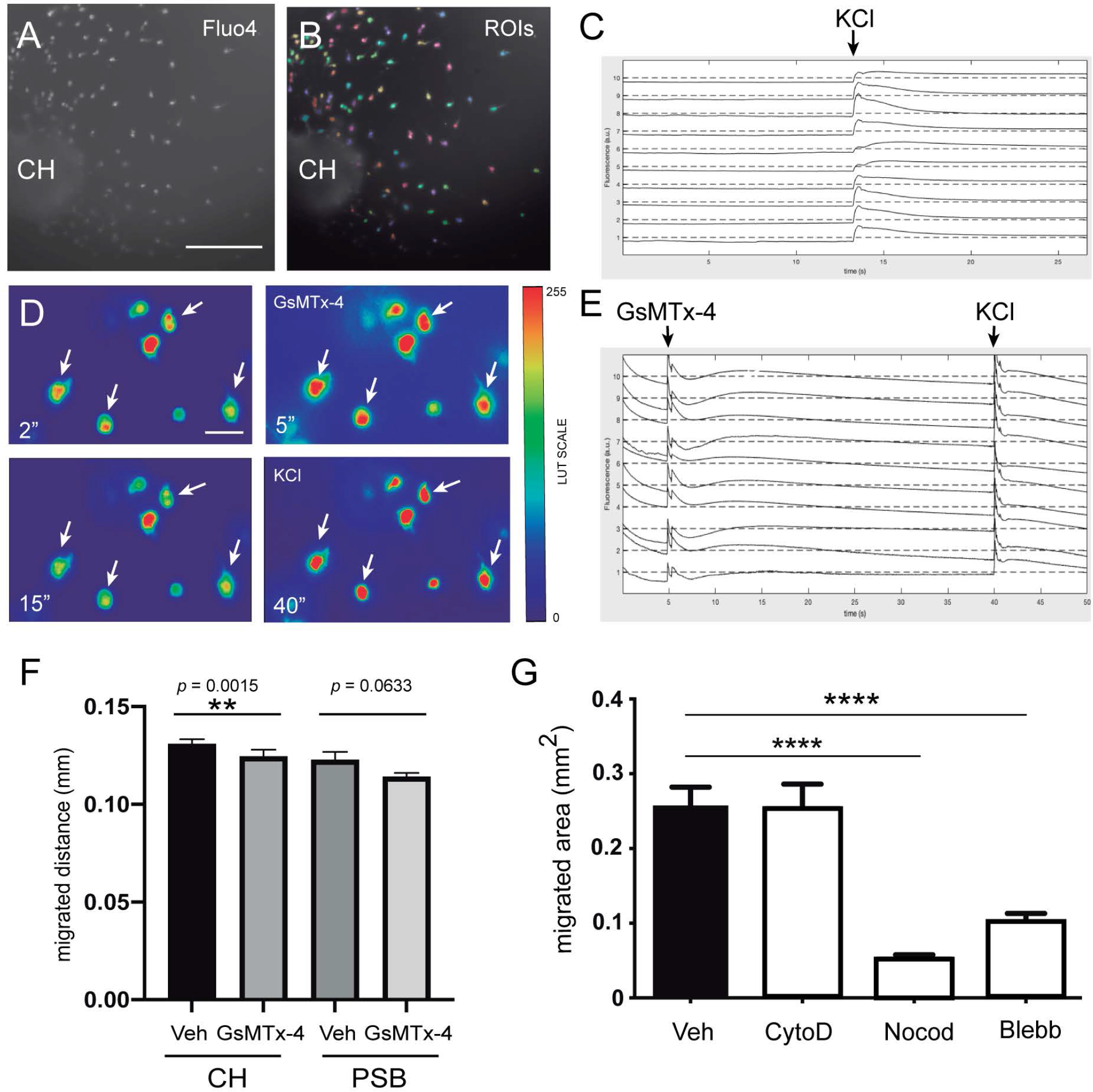
Involvement of mechanosensory receptors in the migration of CH-derived CR cells. **A-C)** Fluo4-AM experiments demonstrating that CR cells are able to depolarize in the presence of KCl. Images in A and B correspond to an example of 2D culture of the CH on methylcellulose-containing medium. During the fluorescence analysis using NetCal^TM^, CR cells were identified and selected by specific individual ROIs (B). **C)** Example of the fluorescence changes of several identified CR cells (y axis). The time (x axis) is that of incubation with 0.1 M KCl (arrow). **D-E)** Frames obtained from identified CR cells illustrating the changes in Ca^2+^ during incubation with GsMTx4 and KCl (see also S3 Movie). A lockup table was applied during movie editing in order to better reveal the transient increase in Ca^2+^ after GsMTx4 (at 5 sec) and KCL (at 40 sec). The changes in the fluorescence ratio after these treatments (labelled as arrows) for several CR cells are illustrated in (E). **F-G)** Histograms illustrating the effects of GsMTx4 treatment on the migration of CH- and PSB-derived CR cells (F) and after Cytochalasin D, Nocodazole, and Blebbistatin treatment (G). Data are presented as mean ± s.e.m. Veh: Vehicle; CytoD: Cytochalasin D; Noco: Nocodazole; Bleb: Blebbistatin; CH: cortical hem; PSB: pallium subpallium boundary. The specific *p* values are included in (F), and ** *p* < 0.05 and **** *p* < 0.0001 in F and G, respectively. Scale bar A = 300 μm pertains to B; D = 50 μm.

### Ectopic transplantation of CH and PSB-derived CR cells demonstrates intrinsic mechanical properties *in vitro*

Due to the above illustrated data, we aimed to develop ectopic transplantation experiments in an *in vitro* preparation of telencephalic slices (see Materials and Methods for details) (Fig. 4). Thus, coronal embryonic telencephalic slices (E12.5) from wild-type mice were cultured on transwells, essentially as described [53], and the endogenous CH and PSB were removed, while the CH and PSB from mTmG reporter mice were transplanted (Fig. 4D). Migrated CR cells generated after explant transplantation could be easily identified by their red fluorescence protein (tdTomato) expression, but also by using double labeling with CALR antibodies. Results demonstrated that CH-derived CR cells transplanted in their original position are able to migrate long distances tangentially in dorsal and medial marginal zones of the slices (mTmG CH in CH location = 565.7 ± 101.3 μm, n = 9) (S2 Fig.). However, PSB-derived CR cells were unable to migrate longer distances in dorsal portions of the pallium (mTmG PSB in CH location = 200.7 ± 72.7 μm, n = 12; all mean ± s.e.m.) (Fig. 4D-E). In contrast, when mTmG CH explants were transplanted in the PSB location, a large number of double-labeled CR cells with mTmG and CALR could migrate dorsally as well as towards ventral portions of the telencephalon (mTmG CH in PSB location, lateral-dorsal migration = 603.1 ± 55.0 μm; lateral-ventral migration = 706.7 ± 79.5 μm; n = 17; all mean ± s.e.m.) (Fig. 4F-K). In parallel, mTmG PSB transplanted in the PSB location showed increased migration when compared after transplantation in the CH (mTmG PSB in PSB, dorsal migration = 310.5 ± 44.25 μm; ventral migration = 232.9 ± 30.94 μm; n = 11; all mean ± s.e.m.) (Fig. 4J-K). From these experiments we may conclude that PSB dorsal migration is *≍* 1.55 times greater when transplanted in PSB than in CH regions. In contrast, CH showed *≍* 1.25 times greater migration distances when transplanted in the PSB than in the CH. These data agree with previous TFM results indicating that CR cells originating from the CH were able to generate stronger mechanical forces to the substrate and they migrate in hydrogels ranking from 5.8 mg/ml to 9.8 mg/ml greater than PSB-derived CR cells. Thus, CH-derived CR cells, when transplanted in PSB, were able to strongly migrate both due to the lesser stiffness of the region (as compared to dorsal regions) and to their intrinsic mechanical properties. In contrast, PSB CR-derived cells migrate less towards or within dorsal pallial regions with increased stiffness. Taken together, the present data suggest that both the differing stiffness (dorsal *vs* lateral) of the developing marginal zone/layer I and the intrinsic differences in the motogenic properties of the CR cells depending on their origin play a role in determining their distribution in the developing marginal zone as observed *in vivo*.

**Fig. 4.**
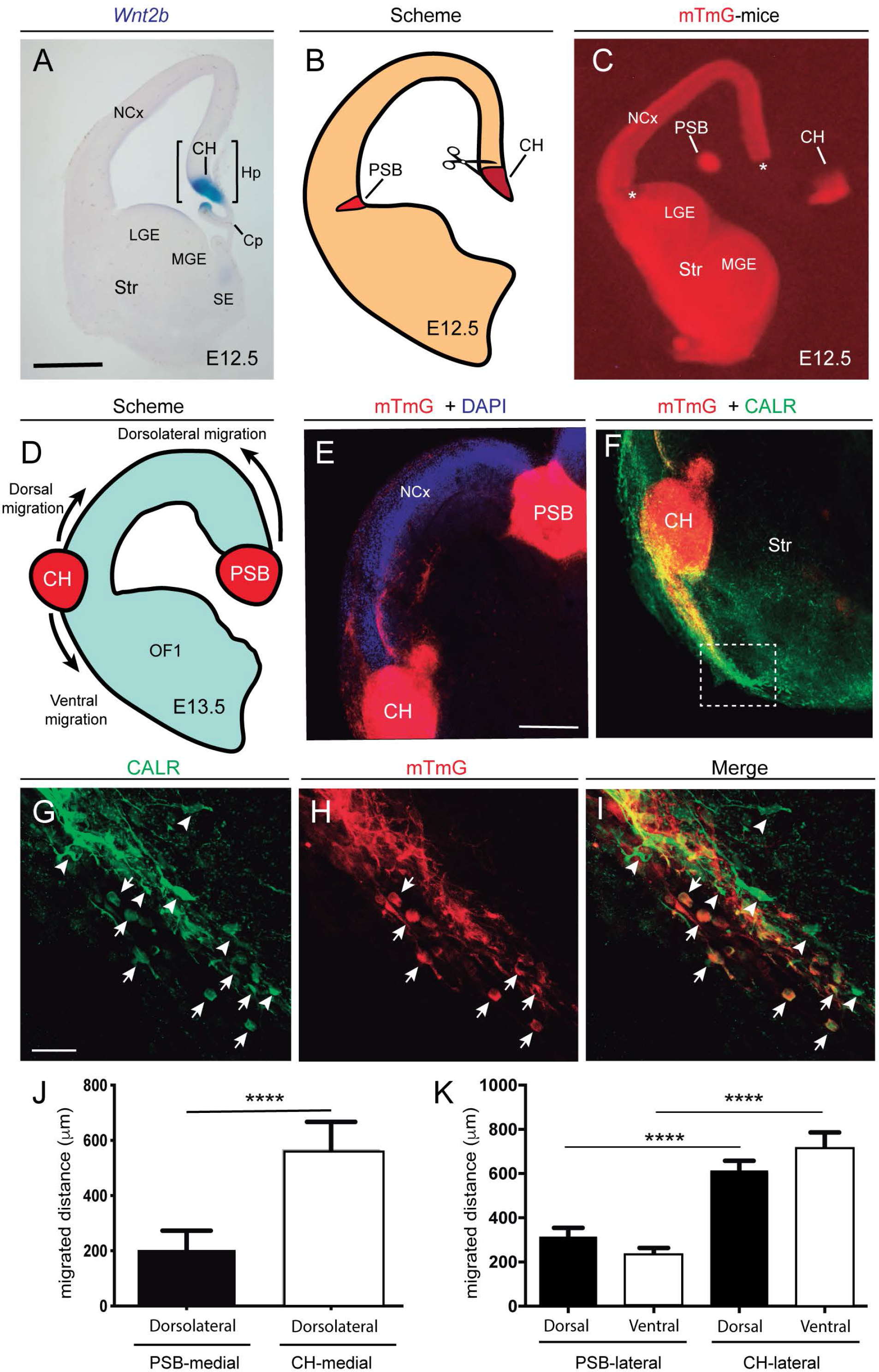
Examples of the differential behavior of CH- and PSB-derived CR cells after transplantation experiments. **A)** Example of an E12.5 coronal section showing the location of the CH after *Wnt2b in situ* hybridization. **B-C)** Scheme (C) and low-power fluorescence photomicrographs (D) illustrating the microdissection procedure for the CH and the PSB using the reporter mTmG mice. **D-E)** Scheme (D) and confocal microcopy photomicrographs (E) illustrating the location of the transplanted CH and PSB in telencephalic slices (see Materials and Methods for details). **F)** Photomicrographs illustrating the migratory stream of CH-derived CR cells after their lateral transplantation. The dashed labelled box is illustrated in G-I. **G-I)** Examples of double-labelled CR cells identified with CALR antibodies and mTmG fluorescence (arrows). Note the large number of double-labelled cells migrating ventrally to the transplanted CH. In contrast, few CALR-positive cells (arrowheads) are mTmG-negative. **J-K)** Histograms illustrating the migrated distance (y axis) of double-labelled (CALR-mTmG) CR cells in the transplantation experiments. In (J) the dorsal-lateral migration of the PSB and CH transplanted in the natural CH location is shown. In (K), the migrated distance of double-labelled cells in the dorsal and ventral regions of the CH and PSB transplants after a lateral transplantation is illustrated. Data are presented as mean ± s.e.m. **** *p* < 0.0001. Abbreviations: Cp = choroid plexus; CH = cortical hem; Hp = hippocampal primordia; LGE and MGE = Lateral and medial ganglionic eminences; NCx = neocortex; SE = septal region; Str = striatum; PSB = pallial subpallial boundary. Scale bars, A = 300 μm pertains to C; E = 200 μm pertains to F and G = 50 μm pertains to F-I.

## Discussion

Several studies in the literature combines lineage analysis, *in vitro* cultures, and CR cell markers, and have revealed the migratory routes of CR cells in developing pallium have been revealed (see among others [16-18, 22, 27, 42, 48, 65, 75, 76]). To summarize these studies, CH-derived CR cells migrate in the marginal zone expanding to dorsal and medial parts of the neocortex with a preponderance in the medial level in the caudal axis (see [68] by way of example), since the more rostral levels are mainly populated by SR-generated CR cells. In contrast, PSB-derived CR cells are more confined to lateral portions of the pallial marginal zone with a lesser presence in the medial and dorsal regions (see [68] by way of example).

Several groups have focused their attention on defining chemical factors that mediate the migration of the different sets of CR cells (see Introduction). Concerning migration properties of subsets of CR cells, an elegant study carried out by Barber *et al.* [65] demonstrated *in vitro* using whole flattened cortical vesicles that CR cells originating in the CH showed increased migratory speed compared to PSB-derived CR cells. The difference (*≍* 18-20-% at E10) was also associated with differences in VAMP3 expression (increased in CH-derived CR cells with respect to PSB-derived ones). VAMP3 is involved in endocytosis, a crucial process that modulates membrane dynamics at the leading edge of migrating neurons and axons [77, 78] as occurs in other cell types [79]. In addition, the authors showed that CH-derived CR cells migrate towards the dorsal cortex while PSB-derived CR cells migrate laterally in the rostro-caudal axis [65]. Our current data corroborate these results.

Concerning expansion of the CR cells in marginal zones, several hypotheses have been proposed. For example, Villar-Cervino *et al.*, indicated that the distribution of different CR-cell subtypes is mediated by a contact repulsion process [65]. These effects were not observed when they analyzed groups of CR cells growing in Matrigel^TM^ [53], nor were they seen in present results. However, Barber *et al.* suggested that, after analysis of their experiments and time-lapse results, factors other than contact repulsion might regulate CR-cell expansion and trajectories in the developing pallium. Our present results are in line with their study, since we show that mechanical factors might contribute to the distribution of CR cells in the dorsal-lateral axis of the developing marginal zone. As indicated, emerging evidence demonstrates that, in parallel to chemical cues, mechanically-mediated processes play relevant roles in axonal guidance, neuronal migration, neurite growth, and brain development [55, 57, 58]. Concerning pallial morphogenesis, recently published studies from the Miyata lab reported new results. In fact, a recent study of Nagasaka *et al*. analyzed the stiffness differences between the ventricular zone and the pallium and ganglionic eminence [80]. Relevantly, in this study the authors reported greater *E* values in the pallial *vs* the subpallial ventricle, which could be a factor involved in pallial folding [80]. Our present results reinforce and expand these observations, demonstrating a significant difference for *E* values between the dorsal part of the marginal zone of the pallium and the lateral regions. However, we cannot rule out the possibility that these differences are also linked to those observed in the ventricular zone. From a developmental point of view, during the early stages of cortical development, the pallium expands in thickness with the addition of postmitotic neurons, generating the cortical plate in a lateral-to-medial gradient, but it also expands laterally [81]. Concerning the lateral-to-ventral expansion of the neocortex, a study by Saito *et al.* described how a migratory stream of the early generated (E10.5) dorsally preplate cells (mainly subplate cells) participated strongly in this lateral expansion, as evidenced by a dorsal-to-lateral migration [24], and most probably acting on the orientation of radial glial cells and generating axon tensions at the level of subpallium, at E14.5 [82]. The presence of these subplate corticofugal axons (labelled with anti-GABA antibodies or 1,1’- Dioctadecyl-3,3,3’,3’-Tetramethylindocarbocyanine Perchlorate (DiI) tracing) at E14.5 was also reported by a number of pioneering studies [83–85]. Under this scenario, the lateral part of the pallium which is closest to the ganglionic eminence will allow the tangential migration of these preplate-derived cells [24]. However, whether these early generated VZ-derived subplate cells with monopolar morphology also show mechanical differences with the non- monopolar cells located in the medial-dorsal portions of the subplate warrants further study and is of interest in fitting their functions and behavior into the stiffness differences observed in the developing pallium.

With respect to CR cells, these studies of Miyata’s lab did not focus on this cell type. Our results demonstrate that CH-derived CR cells can migrate long distances in the marginal zone in both the lateral and the dorsal part of the pallium. Interestingly, they were able to migrate more in the lateral than the dorsal regions in our transplantation experiments. In contrast, PSB-derived CR cells showed reduced migration when transplanted into dorsal regions of the neocortex. This observation is likely related to their migratory properties in the differing stiffness of the dorsal *vs* lateral parts of the marginal zone (BIO-AFM experiments) as corroborated in our Matrigel^TM^ experiments. Taking this into account, our observations and those of Saito *et al.* suggest that the migration of CR cells follows the changes in cortical stiffness generated between E10 and E12.5. This is of relevance since in the absence of these coordinated actions, CR cell mechanical properties, dorsal-lateral *E* differences in the ventricular and marginal zones, as well as the described dorsal-lateral stream of preplate-derived cells, might trigger altered neocortical development. In fact, it is widely recognized that correct CR cell distribution in marginal zone-layer I is a crucial factor in both radial glia maintenance [86] and neuronal radial migration [87]. With altered CR cell distribution, changes in cortical plate development and layer specification might occur (e.g., [22, 86, 88]). Our study describes for the first time how subsets of CR cells can also be characterized by their mechanical properties which, along with the differential dorsal-lateral stiffness of the marginal zone and additional chemical cues, allow the orchestrated dorsal-lateral migration of two different CR-cell populations (CH and PSB-derived) leading to a specific regional distribution that plays a role in cortical development and maturation.

### Coordinated action of mechanical and chemical cues modulates the dorsal-lateral migration and distribution of CH and PSB-derived CR cells: A putative scenario in early neocortical development

In Fig. 5 we hypothesize a putative scenario for the dorsal-lateral CR-cell migration generated in the CH and the PSB. This summary includes results published by several research groups and the present results. In this scheme, the dorsal-medial difference in stiffness is illustrated (present results) and the presence of CXCL12 and Sema3E is illustrated. In addition, we include data related to the expression of CXCR4 as well as PlexinD1. In this hypothesis, during development, SR-, CH-, and PSB-derived CR cells are generated in parallel. However, neither the CH nor the PSB express PlexinD1 or CXCR4 (Fig. 5A-D,F). In contrast, in the marginal zone, both CH- and PSB-derived CR cells express PlexinD1 and CXCR4, as well as Reelin (Fig. 5A-F). Due to the differing stiffness of the pallium, PSB-derived CR cells can migrate in their lateral portions to the marginal zone, while being blocked in dorsal pallial regions displaying greater stiffness and increased Sema3E expression. In contrast, CH cells with greater mechanical properties can migrate in dorsal portions but are also progressively affected by the action of Sema3E (Fig. 5G). Both CR-cell populations are positioned in the marginal zone by the action of CXCL12, and their final distribution in the dorsal-lateral axis is promoted by the inhibition of CXCL12/CXCR4 signaling by Sema3E, as demonstrated in [53], along with their differences in intrinsic mechanical properties and the stiffness of the developing pallium (present results). In addition to this, other factors such as the expression of VAMP3 leading to intrinsic differences in CR-cell migration as well as other repulsive actions described in other studies play crucial roles in parallel to accomplish their regional distribution in the rostral-caudal and medial-lateral axis of the pallium (see Introduction for details).

**Fig. 5.**
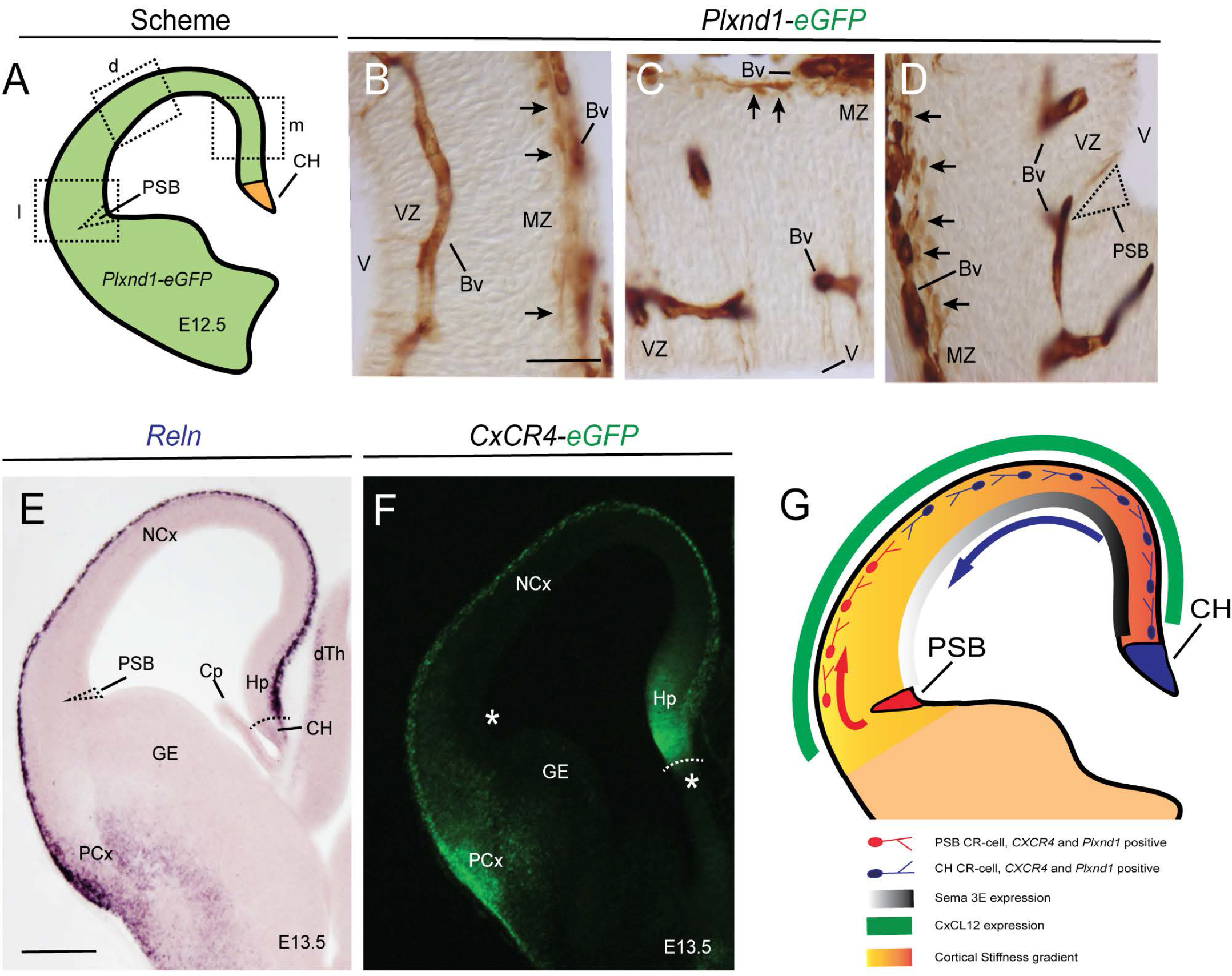
Mechanical and chemical cues modulate the migration of PSB- and CH-derived CR cells. Data presented in this figure illustrate some of the cues involved in the CR-cell migration. **A-D)** Scheme (A) and eGFP immunostaining using immunoperoxidase methods of PlexinD1 in transgenic *Plxnd1*-eGFP mice. PlexinD1 is present in blood vessels as well as in CR cells in the complete marginal zone (arrows in B-D). The specific location of the PSB is labelled in D illustrating the absence of PlexinD1 in this region and demonstrating that its expression is postmitotic and linked to its position in marginal zone. **E-G)** Reln *in situ* hybridization (E) and eGFP fluorescence in CxCR4-eGFP mice (F), respectively, illustrating their distribution in the developing pallium. Asterisks and dashed regions in F and G illustrate the absence of *Reln* and CXCR4 in the proliferative regions, especially the PSB. **G)** Scheme summarizing the results observed in our studies (see Discussion for details). Abbreviations as in Fig. 4 in addition to Bv = blood vessel; dTh = dorsal thalamus; GE = ganglionic eminence; MZ = marginal zone; PCx = piriform cortex; V = lateral ventricle; VZ = ventricular zone. Scale bars, B = 50 μm pertains to C-D; E = 300 μm pertains to F.

## Material and Methods

### Animals

The following mice and rat strains were used in the present study: OF1 mice (E12.5) and Sprague Dawley rats (E14.5) were purchased from Charles River laboratories (Paris, France). In addition, the mTmG mice (ROSA^mT/mG^; Stock No. 007576; The Jackson Laboratories; Bar Harbor, ME, USA) were also used. *Plxnd1-eGFP* mice were obtained from the Mutant Mouse Regional Resource Center (MMRRC; University of California, CA, USA). mTmG and *Plxnd1-eGFP* mouse genotypes were verified by observing a tail fragment under a fluorescence microscope (BX61; Olympus Corporation). In addition, CXCR4-eGFP transgenic mice (kindly provided by J.H.R. Lubke (Germany) [89]) were used. All the animals were kept in the animal facility of the Faculty of Pharmacy at the University of Barcelona under controlled environmental conditions and were provided food and drink *ad libitum*. For AFM experiments, OF1 pregnant mice were housed in the animal facility of the Faculty of Medicine at the University of Barcelona. All the experiments were carried out following protocols of the Ethics Committee for Animal Experimentation (CEEA) of the University of Barcelona (OB47/19, C-007, 276/16 and 47/20).

### CH and PSB explant dissection

In order to obtain CH and PSB regions, brain embryos (E12.5) were covered with a mixture of L15 medium (31415-029, Invitrogen) containing 4% low melting point agarose (50111, Lonza) which was then allowed to solidify at 4°C for a few minutes. After gelation 300 µm-thick coronal sections were obtained using a vibratome (Leica). Free-floating sections were collected in cold 0.1M PBS containing 0.6% glucose and selected under a dissection microscope, and the explants of CH or PSB were dissected.

### Culture of CH and PSB explants on hydrogels

For explants embedded in hydrogel, a sandwich procedure was developed using a base of homemade rat tail collagen [90] and a top of Matrigel^TM^ (354434, Corning, Cultek). To prepare the Matrigel^TM^ at different densities, cold Neurobasal^TM^ media (21103-049, Invitrogen) was used to dilute Matrigel^TM^ stock solution. One explant per preparation was placed on a homogeneous collagen base and then another layer of Matrigel^TM^ was added on top. Before the Matrigel^TM^ gels, explant position was verified and correctly positioned under the dissecting microscope. Once the explant was seeded, the gel was allowed to coagulate at 37°C, 5% CO2 before adding the supplemented medium. At 2 days in vitro (DIV), explants were fixed for 1 hour with cold 4% buffered paraformaldehyde (PFA) before washing with 0.1 M PBS, and then stored at 4°C prior to immunocytochemistry or photodocumentation. The number of cells that migrated out of the explants was counted, and the maximum distance migrated was also determined using FIJI^TM^ software, using as calibration pictures a millimetric eyepiece at the same magnification. CH-derived explants were treated with 2 μg/ml of Cytochalasin D (C8273, Sigma-Aldrich), 10 μM Nocodazole (M1404, Sigma-Aldrich), or 0.5 μg/ml of Blebbistatin (1760, Tocris).

### *In vitro* transplantation of CH or PSB explants in telencephalic slices

Brain slices were obtained from E12.5 wild-type mice and the CH and the PSB from E12.5 mTmG mouse embryos as described above (see also [53]). Slices were transferred to collagen-coated culture membrane (PICM0RG50, Millipore) in 1.2 ml of medium BME-F12 1:1 (41010-026, Invitrogen), glutamine (25030-024, Invitrogen), 5% horse serum (26050-088; Invitrogen), penicillin, streptomycin (15140122, Invitrogen), and 5% bovine calf serum (12133C, Sigma-Aldrich). The CH and PSB from wild-type slices were discarded and replaced with the dissected CH or the PSB from mTmG mice slices. After several hours, the medium was changed to Neurobasal^TM^ medium supplemented as above and cultured for up to 48 hours before analysis.

### *In situ* hybridization

*In situ* hybridization was carried out as described previously [91, 92] on 50 μm vibratome fixed brain sections of E12.5 embryos. Both sense and antisense riboprobes against *Wnt2b* (provided by P. Bovolenta) and *Reln* (provided by T. Curran) were labelled with digoxigenin, according to the manufacturer’s instructions (Roche Farma).

### Immunocytochemical methods

The processing of each *in vitro* model was determined by its culture characteristics (hydrogel, coverslip, or brain slice). Although the general procedure was similar for all conditions, the incubation times and mounting methods for analysis differed. The general procedure started with fixing the tissue samples with 4% PFA, then washing them with 0.1M PBS and a blocking solution composed of 10% Fetal Bovine Serum (FBS; 10500064, Invitrogen) and 0.1M PBS with Triton X-100 (X100, Sigma-Aldrich) (concentration determined by each model). After washing with PBS-Triton X-100, the primary antibody was incubated with 7% FBS and PBS-0.2% gelatin and Triton X-100. This was followed by the Alexa-tagged secondary antibodies (Alexa 488; A21206, Invitrogen) diluted in 7% serum (FBS) and PBS 0.1M containing gelatin 0.2% and 0.1 % Triton X-100. Afterwards, nuclear staining was performed using Hoechst (1 µg/ml; B2261, Sigma-Aldrich). The antibody used for CR-cell labeling was Calretinin (CALR; 1:1000; 7697, Swant). Details of the immunocytochemical procedures in each model are briefly explained below. For explant cultures growing in Matrigel^TM^ 0.5% Triton X-100 was used in all steps and long incubation times were observed. After fixation for 1h at 4°C with 4% buffered PFA and blockade for 4h at room temperature, the primary antibody was incubated for 2 overnights at 4°C and the secondary antibody for 1 overnight at 4°C with gentle shaking. Finally, nuclear labeling with Hoechst was developed for 20 minutes before washing with 0.1M PBS and mounting with Mowiol^TM^ (475904, Calbiochem). For primary cultures of CH or PSB CR cells growing on (PAA) gels for traction force microscopy (TFM) experiments, a short fixation time (5 min) with 2% buffered PFA (removing half of the medium), and 10 minutes in 4% buffered PFA at 4°C, was developed in selected preparations. Thereafter, fixed neurons were incubated at room temperature for 1h with blocking solution, 2h with primary antibodies, and 1h for the secondary antibodies, at room temperature with gentle shaking. Hoechst staining was run for 10 minutes before washing with 0.1M PBS after immunohistochemistry and photodocumentation. For slice cultures, coronal slices after CH or PSB transplantation (see above) were fixed for 1 h with 4% buffered PFA, and after this detached from the transwell membrane and processed in flotation and gentle agitation. All the immunohistological solutions contained 0.5% Triton X-100 and the samples were incubated for longer times. Thus, slices were treated with blocking solution for 4h, incubated with primary antibodies for 48 hours, and then for 12 hours with secondary antibodies at 4°C with gentle shaking. Finally, the Hoechst solution was incubated for 20 minutes before washing with 0.1M PBS and mounting with Mowiol^TM^; double CALR- mTmG-positive CR cells were photodocumented using a Zeiss SLM400 confocal microscopy.

### Viscoelastic properties of Matrigel^TM^ hydrogels analysed with rheometry

The different densities checked in the study were obtained from a Matrigel^TM^ stock with known total protein concentration diluted in cold Neurobasal^TM^ medium. The dilutions were different for different batches of Matrigel^TM^: 12.72 mg/ml (Lot n°.: 9294006), 12.6 mg/ml (Lot n°: 8015325), 11.95 mg/ml (Lot n°: 9021221), and 9.8 mg/ml (Lot n°: 9148009). To obtain the hydrogels, a minimum of 100 µl of mixture was needed per dish, carefully depositing the mixture and leaving it to gel for 2-4 hours at 37°C. Once gelled, complete Neurobasal^TM^ medium was added to cover, and left in an incubator at 37°C and 5% CO2. At 2 DIV the dishes were carefully lifted and placed on a rheometer plate previously heated to 37°C and calibrated. For rheometry measurement of the hydrogels, an *Ø* 8mm Peltier (Peltier plate Steel - 108990) coupled to a Discovery Hybrid Rheometer HR-2 (Discovery HR-2; 5332-0316; TA instruments) was used. To prevent evaporation, rapeseed oil was used as a solvent-trap after applying loading gap to the sample and deleting medium and excess hydrogel. Finally, the geometry was taken to the gap to geometry and the chosen measurement began. With the TRIOS program (v. 5.0.0.44608) the Frequency sweep test using a gap of 0.5 mm (previously determined) was selected. To obtain the elastic modulus *E*, we first measured the storage and loss moduli in experiments at 1 Hz frequency sweep, which provided a strain modulus G given by the equation (1).

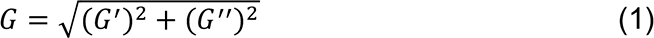

Where *G’* corresponds to storage modulus and *G’’* to loss modulus. Once *G* was obtained, the elastic modulus *E* values were determined as equation (2).

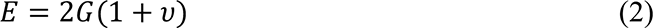

Where *V* is the Poisson’s ratio, defined as the ratio of transverse contraction strain to longitudinal extension strain, and that was assumed to be 0.5 for low stiffness hydrogels.

### Atomic force microscopy (AFM) experiments

For *in situ* AFM measurements, E12.5 mouse embryonic brains were carefully dissected without damaging the cortical surface. Then, a 4% agarose (SeaPlaque^TM^ GTGTM Agarose, 50111; Lonza) solution was prepared in 0.1M PBS and left at 45°C in a dry bath. The two mouse hemispheres were dissected and examined for dorsal and lateral AFM analysis (Fig. 1). Two brain orientations were generated in embedding telencephalic regions with agarose to achieve dorsal and lateral AFM measurements of the upper surface of the brain (Fig. 1B). Briefly, a large plate, with a glass slide for AFM calibration, containing 2-3 mm thickness of 4% agarose, was prepared. Once solidified, one hemisphere was carefully placed over the agarose surface adding more agarose to embed the ventral subpallial brain regions, leaving the dorsal pallial region to develop the dorsal AFM measurements (Fig. 1B). In parallel experiments, the other hemisphere was placed laterally over the bottom agarose with the medial brain portion in contact with agarose, being leaving the lateral part of the agarose-embedded brain for AFM measurement (Fig. 1B). After gelling, the plate was covered with complete Neurobasal^TM^ medium and placed in the incubation chamber at 37°C.

Measurements were carried out on a custom-made BIO-AFM mounted on an inverted optical microscope (TE2000; Nikon). AFM was equipped with a V-shaped silicon nitride cantilever (0.01 N/m nominal spring constant) terminating in a 6 μm-radius borosilicate spherical tip (Novascan Technologies). The cantilever deflection was measured by using the optical lever method, and the sensitivity of the photodiode was calibrated prior to probing each sample by using the agarose semi-embedded glass slide in the preparation as reference. For each measurement (dorsal or lateral), 4 separate probing points were selected by laterally displacing the AFM probe 40 µm between measurements. For each probing point, the elastic modulus was calculated from the force-displacement curves by adjusting the Hertz model for the tip-surface contact [93]. From these 4 separate values, the average was calculated, and data were represented by mean and standard error of the mean (s.e.m.) for each brain at the different positions.

### Primary cultures and TFM measurements of CH or PSB-derived CR cells

Pieces of CH and PSB were obtained as above and collected in cold dissection media (0.1M PBS (14200, Invitrogen) and 0.65% glucose (G8769, Sigma-Aldrich)) and centrifuged for 5 minutes at 800 rpm. After removal of the dissection media, 3 ml of dissection medium containing 10X trypsin (15400-054, Invitrogen) at 37°C for 15 minutes was added. After digestion and inactivation with heat-inactivated normal horse serum (1:3 ratio), 10X DNase (AM2222, Ambion) diluted in fresh dissection media was added, and incubated for 15 minutes at 37°C. Finally, 10 ml of dissection medium was added and centrifuged for 5 minutes at 800 rpm. The pellet was resuspended in complete Neurobasal^TM^ medium, and the cells were seeded on the plate. Then cell density was approximately 100,000 cells per 9.5 cm^2^ plate. For acrylamide gels, a Matrigel^TM^ coating (diluted 1:40) was made the day before and incubated at 37°C overnight. The next day, the surface was rinsed with Neurobasal^TM^ medium. For the TFM assay, only isolated cells with clear morphology of CR cells (see below) were analysed to avoid interference from traction forces between different cells. PAA gels with different stiffness were generated by modifying the proportion of acrylamide 40% (1610140; BioRad) and Bis-acrylamide Solution (2% w/v; 10193523; ThermoFisher) and the level of crosslinking in the gel (between *≍* 40 Pa up to supraphysiological values) [94]. To detect gel displacements due to CR-cell mediated forces, the PAA gels was labelled with FluoSpheres® Carboxylate-Modified Microspheres 0.2 µm ((625/645) F8806; LifeTechnologies). In order to generate a good homogeneous Matrigel^TM^ coating for CR cells, the PAA gels were treated with SulfoSANPAH (803332; Sigma-Aldrich). Thus, 0.1M PBS was removed and a mixture of SulfoSANPAH (and 20 µl of reagent diluted in 480 µl bidistilled water) was added. After two rinses with 0.1M PBS, treated gels were coated overnight at 37°C. The following day, coated gels were washed with the fresh culture medium and allowed to stabilize for a few minutes before seeding with CR cells. Once adhered, isolated CR cells were selected (40x objective, inverted Olympus microscope IX71) and placed on top of the PAA gel. For image acquisition and data processing, a Matlab^TM^ script was used (see [95] for details), and the displacements were represented on the bright field image of the cell.

### Calcium analysis in cultured CR cells with Fluo4-AM

To develop analysis of the changes in Ca^2+^ levels in CR cells, CH-derived explants were cultured on Matrigel^TM^ coated dishes as indicated above. In order to enhance explant adhesion and correct CR migration, culture media contained 1% methylcellulose. In previous experiments, we determined the appropriate concentration of methylcellulose and the morphology of CALR-positive cells (S1 Fig.). After 24 hours of culturing, explants were incubated for 30 min with the cell– permeant calcium-sensitive dye Fluo4-AM (F14201, Molecular Probes). The culture was washed with fresh medium after incubation and finally placed in a recording chamber for observation. The recording chamber was mounted on an Olympus inverted microscope equipped with a Hamamatsu Orca Flash 4.0 CMOS camera (Hamamatsu Photonics). Cultures were recorded and images (1024 x 1024 pixels) were captured using a 20x objective and 470 nm wavelength (CoolLED’s pE-300^white^, Delta Optics) every 50 ms for 1 min using the CellSens^TM^ software (Olympus). The recordings were analyzed offline using the Matlab^TM^ toolbox NETCAL (www.itsnetcal.com). Identified CR cells were associated with a single region of interest (ROI). The average fluorescence *Fi (t)* in each ROI (CR cell) *i* along the recording was then extracted, corrected for global drifts and artifacts, and finally normalized as *(Fi (t) - F(0,i)) / F(0,i) = fi (t)*, where *F0,i* is the background fluorescence of the ROI. The time series of *fi (t)* was analyzed with NETCAL to determine sharp calcium transients and that reveal neuronal activity. Obtained movies were edited in Fiji^TM^ and the lockup table ’physics‘ was applied. In this experiment, the mechanosensory channel inhibitor GsMTx-4 (ab141871, Abcam) was used at a final concentration of 10 μg/ml during video recording.

### Statistical analysis

Data in this manuscript are expressed as mean ± s.e.m. of at least four independent experiments unless specified. Means were compared using the Mann-Whitney *U* non-parametric test. The asterisks *, ** and *** indicate *p* < 0.05, *p* < 0.01, and *p* < 0.001, respectively. For TFM and AFM analysis, a permutation test (one tail) was performed and *, ** indicate *p* < 0.05 and *p* < 0.01, respectively was considered statistically significant. Statistical test and graphical representation was performed with Prism v.8 (GraphPad Software), RStudio (RStudio, PBC) and R software (The R Foundation).

## Acknowledgments

The authors thank Tom Yohannan for editorial advice and all members of the Del Río, Navajas, Farré, Trepat, Ferrer, Soriano and Ortín labs for their comments. JADR was supported by *PRPSEM* Project with ref. RTI2018-099773-B-I00 from MCINN/AEI/10.13039/501100011033/ FEDER “Una manera de hacer Europa”, the CERCA Programme, and the Generalitat de Catalunya (SGR2017-648). The project leading to these results also received funding from “la Caixa” Foundation (ID 100010434) under the agreement LCF/PR/HR19/52160007 and the María de Maeztu Unit of Excellence (Institute of Neurosciences, University of Barcelona) MDM-2017-0729, to JADR. JS was funded by Generalitat de Catalunya (SGR- 2017-1061) and MCINN/AEI/10.13039/501100011033/ (PID2019-108842GB-C21). XT was funded by Generalitat de Catalunya (SGR-2017-01602), Spanish Ministry for Science and Innovation MCINN/AEI/10.13039/501100011033/FEDER (PGC2018-099645-B-I00), European Research Council (Adv-883739), Fundació la Marató de TV3 (project 201903-30-31-32) and La Caixa Foundation (LCF/PR/HR20/52400004). DN was financed by H2020 European Research and Innovation Programme under the Marie Skłodowska-Curie grant agreement “Phys2BioMed” contract no. 812772. RS was funded by MCINN/AEI/10.13039/501100011033/ FEDER (RTI2018-101256-J-I00). AL-M. was supported by “La Caixa” Foundation and by FPI Programme (BES-2016-076893) supported by Spanish Ministry of Science and Innovation. FMV was supported by FPU Programme (16/03992), from the Spanish Ministry of Universities. The funders had no role in study design, data collection and analysis, decision to publish, or preparation of the manuscript.

