## Supplementary information for "Involvement of mechanical cues in the migration of Cajal-Retzius cells in the marginal zone during neocortical development"

**S1 Fig Characterization of the explant-derived CR cells growing in 2D cultures with different percentages of methylcellulose**. **A-B)** Histograms illustrating the percentage of explant attachment (A) and CR cells' migrated distance (B) with a difference in the percentage of methylcellulose in the culture media. C-D) Low and high magnification of CR cells growing in 1% methylcellulose medium after CALR immunostaining showing their monopolar morphology. Data are presented as mean ± s.e.m. **** $p < 0.0001$. Scale bars in C = 150 μm and D = 75 μm.

**S2 Fig. mTmG-positive CH-derived CR cells migration in the medial wall of telencephalic slice A-B)** Confocal microscopy images illustrating the migration of the mTmG-positive CR cells after CH explant transplantation in the original location (see Material and Methods for details) of E13.5 OF1 slices. Note the great migration distance of mTmG-positive cells (arrows in (B)) in the dorsal and lateral parts of the marginal zone (MZ) of the cultured OF1 slice. Scale bars in C = 200 μm and D = 50 μm.

**S1-2 Movies**: Examples of the differential migration between CH and PSB-derived CR cells in different concentrations of Matrigel$^{TM}$ (7.8 and 9.8 mg/ml). Movie frame rate 25 fps. Each frame was obtained after 8 minutes in culture.

**S3 Movie.** Example of the intracellular $Ca^{2+}$ changes in CR cells derived from the CH using Fluo4-AM. For review purposes the movie frame rate was modified at 100 fps. A frame was obtained every 100 ms. A lockup scale (physics) was applied in Fiji$^{TM}$ ranking from blue (0 grey level) up to red (255 grey level). With the present frame rate, in the movie GsTMx-4 was applied at 2 sec and the KCl at 17 sec of the movie. Selected frames of this movie are included in Fig. 2.
