## Supplementary figures and images for "Involvement of mechanical cues in the migration of Cajal-Retzius cells in the marginal zone during neocortical development"

### Supplementary Figure 1

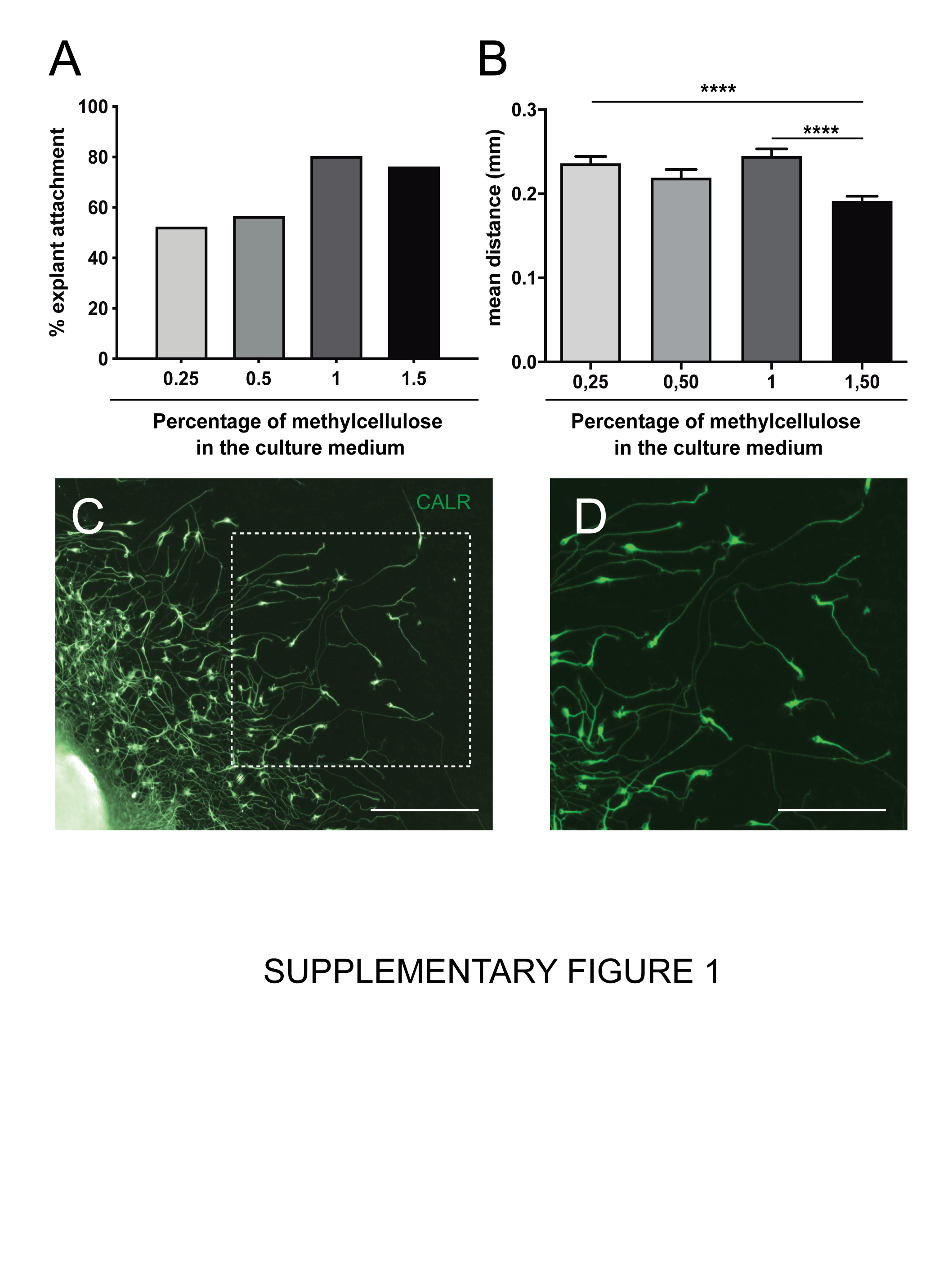

### Supplementary Figure 2

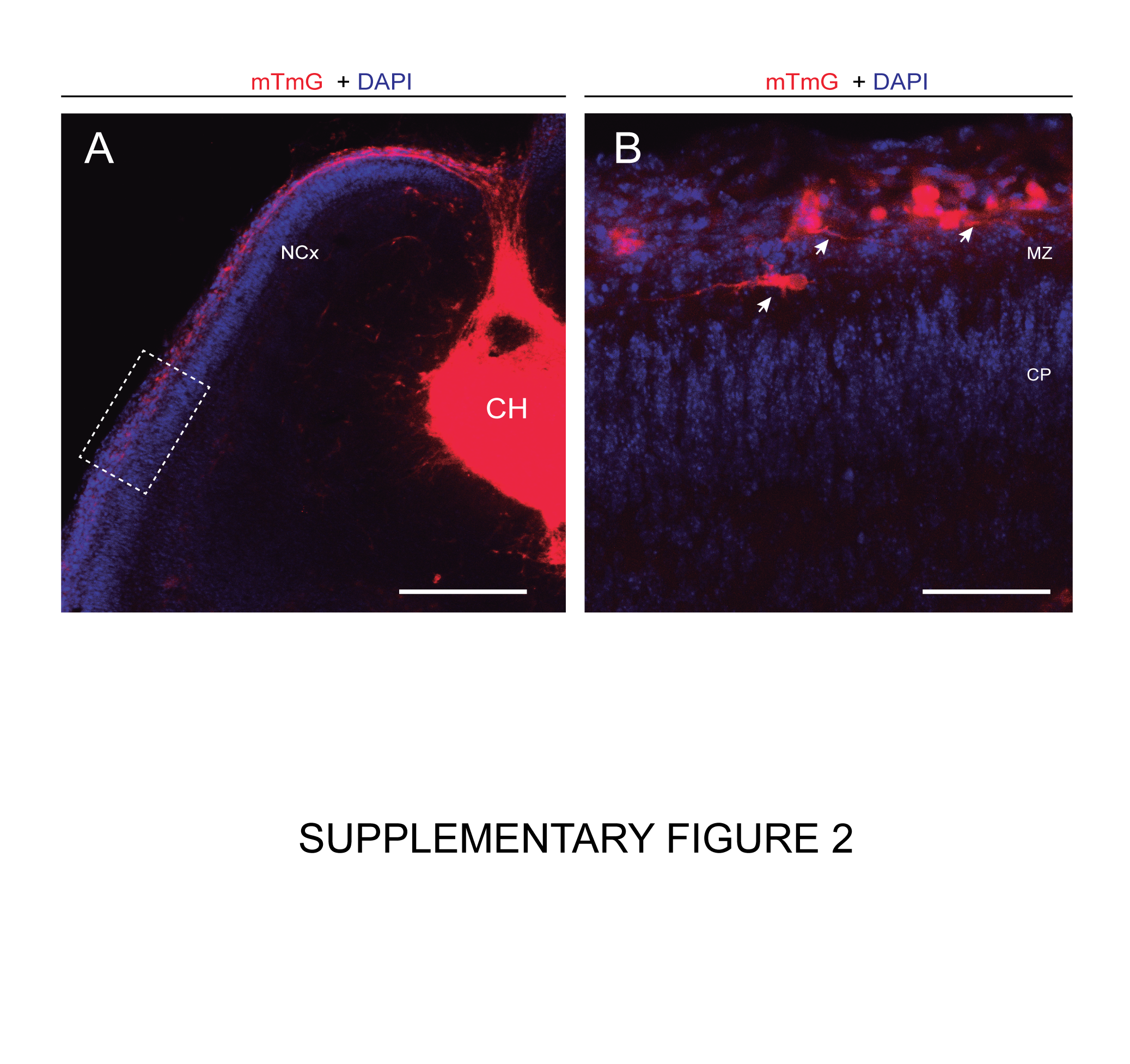
